# Structure of replicating SARS-CoV-2 polymerase

**DOI:** 10.1101/2020.04.27.063180

**Authors:** Hauke S. Hillen, Goran Kokic, Lucas Farnung, Christian Dienemann, Dimitry Tegunov, Patrick Cramer

**Author notes:** These authors contributed equally (Hauke S. Hillen, Goran Kokic, Lucas Farnung, Christian Dienemann, Dimitry Tegunov).

## Abstract

The coronavirus SARS-CoV-2 uses an RNA-dependent RNA polymerase (RdRp) for the replication of its genome and the transcription of its genes. Here we present the cryo-electron microscopic structure of the SARS-CoV-2 RdRp in its replicating form. The structure comprises the viral proteins nsp12, nsp8, and nsp7, and over two turns of RNA template-product duplex. The active site cleft of nsp12 binds the first turn of RNA and mediates RdRp activity with conserved residues. Two copies of nsp8 bind to opposite sides of the cleft and position the RNA duplex as it exits. Long helical extensions in nsp8 protrude along exiting RNA, forming positively charged ‘sliding poles’ that may enable processive replication of the long coronavirus genome. Our results will allow for a detailed analysis of the inhibitory mechanisms used by antivirals such as remdesivir, which is currently in clinical trials for the treatment of coronavirus disease 2019 (COVID-19).

---

Coronaviruses are positive-strand RNA viruses that pose a major health risk^1^. The novel severe acute respiratory syndrome coronavirus-2 (SARS-CoV-2)^2,3^ has caused a pandemic referred to as coronavirus disease 2019 (COVID-19). Coronaviruses use a RNA-dependent RNA polymerase (RdRp) complex for the replication of their genome, and for sub-genomic transcription of their genes^4,5^. This RdRp complex is the target for antiviral nucleoside analogue inhibitors, such as remdesivir^6,7^. Remdesivir shows antiviral activity against coronaviruses in cell culture and animals^8^, inhibits coronavirus RdRp^9,10^, and is currently tested in the clinic as a drug candidate for treating COVID-19 patients^11–13^.

The RdRp of SARS-CoV-2 is composed of a catalytic subunit called non-structural protein (nsp) 12^14^, and two accessory subunits, nsp8 and nsp7^5,15^. The structure of the RdRp was recently reported^16^ and is highly similar to the RdRp of SARS-CoV^17^, a zoonotic coronavirus that spread into the human population in 2002^1^. The nsp12 subunit contains an N-terminal nidovirus RdRp-associated nucleotidyltransferase (NiRAN) domain, an interface domain, and a C-terminal RdRp domain^16,17^. The RdRp domain resembles a right hand and comprises the fingers, palm, and thumb subdomains^16,17^ that are found in all single-subunit polymerases. Subunit nsp7 binds to the thumb, whereas two copies of nsp8 bind to the fingers and thumb sub-domains^16,17^. Structural information is also available for isolated nsp8-nsp7 complexes^18,19^.

To obtain the structure of the SARS-CoV-2 RdRp in its active form, we prepared recombinant nsp12, nsp8 and nsp7 (**Fig. 1a, Experimental procedures**). When added to a minimal RNA substrate (**Fig. 1b**), the purified proteins gave rise to RNA-dependent RNA extension activity, which depended on the presence of nsp8 and nsp7 (**Fig. 1c**). We assembled and purified a stable RdRp-RNA complex and collected single-particle cryo-electron microscopy (cryo-EM) data (**Extended Data Figure 1, Extended Data Table 1**). Particle classification yielded a 3D reconstruction at a nominal resolution of 2.9 Å and led to a refined structure of the RdRp-RNA complex (**Extended Data Figures 1 and 2**).

**Figure 1.**
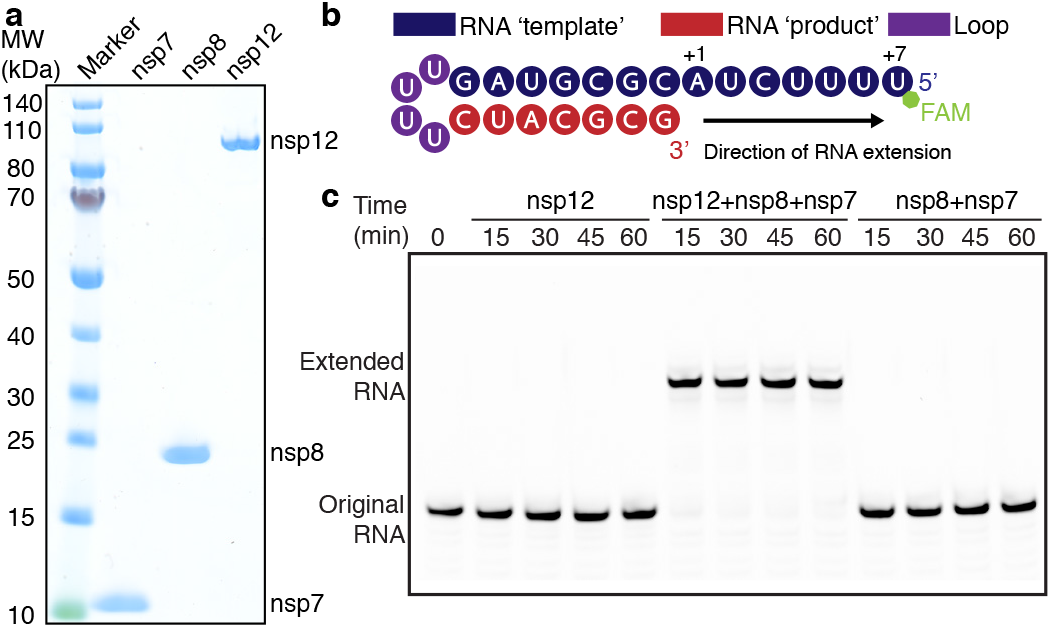
Preparation of active SARS-CoV-2 RdRp. **a** SDS-PAGE analysis of purified SARS-CoV-2 RdRp subunits nsp12, nsp8 and nsp7. **b** Minimal RNA substrate that folds into a hairpin with ‘template’ and ‘product’ regions. The RNA contains a 7-nucleotide fluorescently labeled 5’-overhang. **c** Incubation of the RdRp subunits (a) with RNA (b) leads to efficient RNA extension. RNAs were separated on a denaturing acrylamide gel and visualized with a Typhoon 95000 FLA Imager.

The structure shows the RdRp enzyme engaged with over two turns of duplex RNA (**Fig. 2**). The structure resembles that of the free enzyme^16^, but also reveals large additional protein regions in nsp8 that became ordered upon RNA binding and interact with RNA far outside the core enzyme (**Extended Data Figure 3a**). These observations are unique, as RdRp complexes of hepatitis C virus^20^, poliovirus^21^, and norovirus^22^ contain only one turn of RNA that is however oriented in a similar way (**Extended Data Figure 3b**).

**Figure 2.**
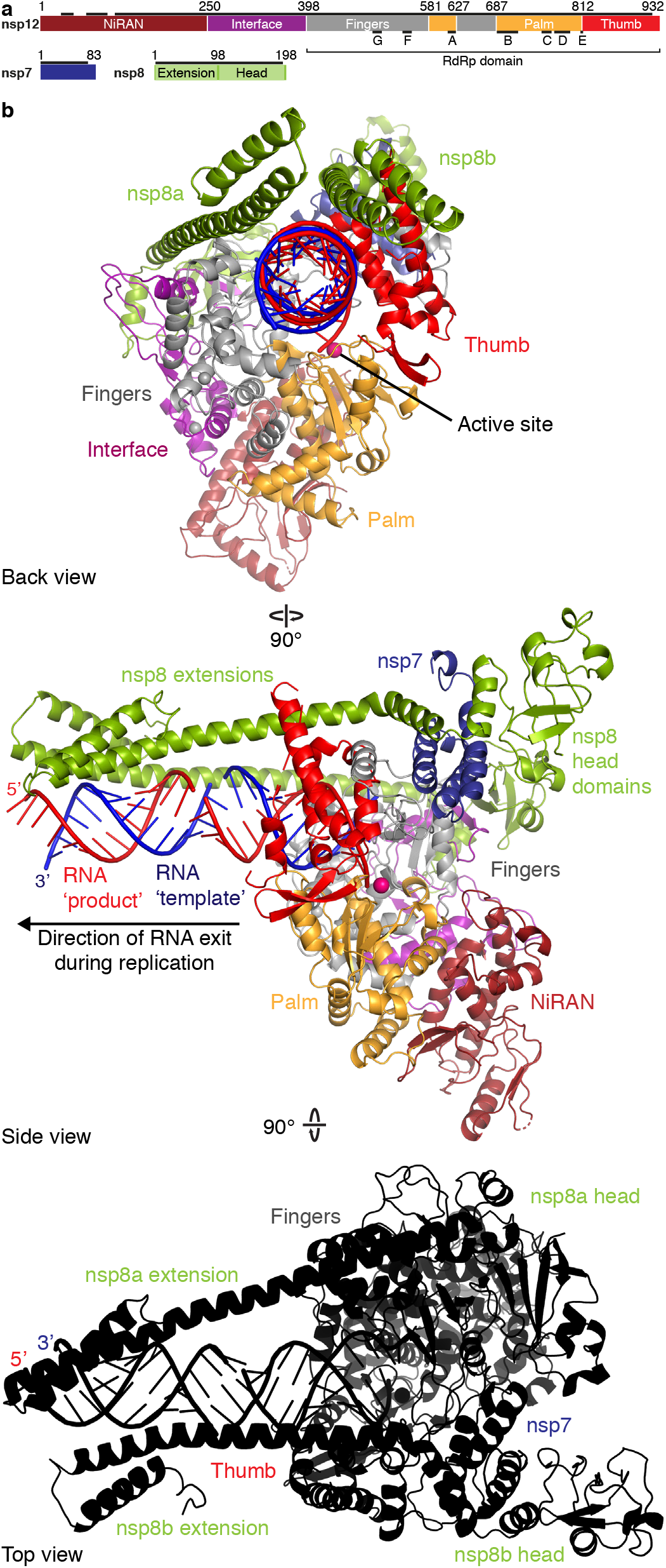
Structure of SARS-CoV-2 RdRp-RNA complex. **a** Domain structure of RdRp subunits nsp12, nsp8, and nsp7. In nsp12, the conserved sequence motifs A-G17 are depicted. Regions included in the structure are indicated with black bars. **b** Three views of the structure, related by 90-degree rotations. Color code for nsp12 (NiRNA, interface, fingers, palm, thumb), nsp8, nsp7, RNA template (blue) and RNA product (red) used throughout. The magenta sphere depicts a modeled^22^ metal ion in the active site.

Our structure shows details of the RdRp-RNA interactions (**Fig. 3**). The nsp12 subunit binds one turn of RNA between its fingers and thumb subdomains (**Fig. 3a, b**). The active site is located on the palm subdomain and formed by five conserved nsp12 elements called motifs A-E (**Fig. 3b**). Motif C binds the RNA 3’-end and contains the residues D760 and D761, of which D760 is known to be essential for RNA synthesis^15^. The additional nsp12 motifs F and G reside in the fingers subdomain and position the RNA template (**Fig. 3b**). The observed nsp12 contacts to the RNA product strand may retain a short RNA during de novo synthesis^15^.

**Figure 3.**
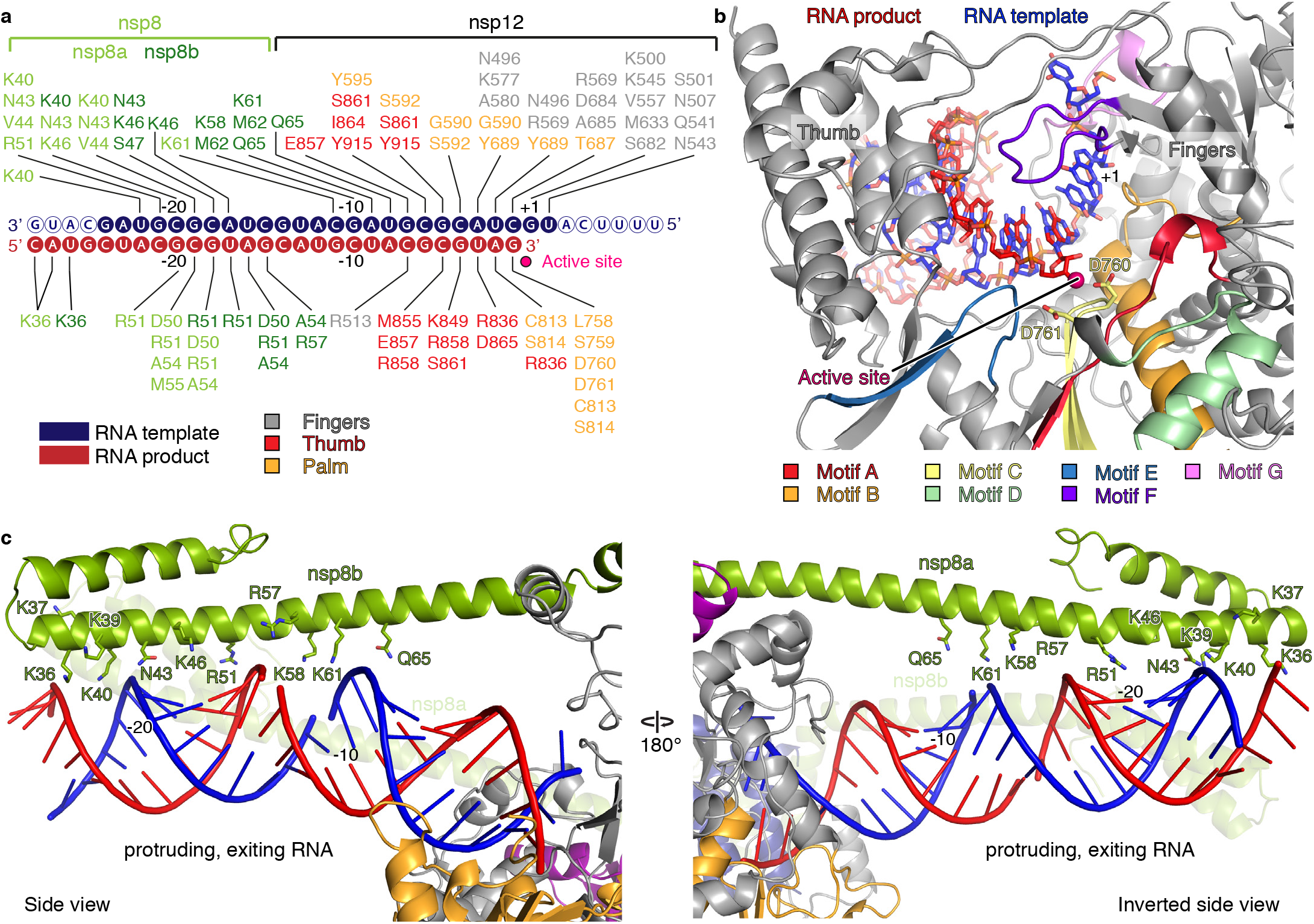
RdRp-RNA interactions. **a** Schematic of protein-RNA interactions. Solid and hollow circles show RNA nucleotides that were included in the structure or not visible, respectively. RdRp residues in nsp12 and nsp8 within 4 Å of the RNA are depicted and contacts indicated with lines. Color code as in Fig. 2. **b** Interactions of RdRp active site with the first turn of RNA. Subunit nsp12 is in grey and conserved motifs A-G are colored as indicated. Active site residues D760 and D761 are shown as sticks. The active site metal ion was modeled^22^ and is shown as a magenta sphere. **c** Interaction of nsp8 subunits with protruding RNA, showing residues in proximity to RNA.

As the RNA duplex exits from the RdRp cleft, there are no structural elements that restrict its extension, consistent with the production of a double-stranded RNA during replication (Fig. 3c). The protruding RNA duplex is flanked by long α-helical extensions that are formed by the highly conserved^18^ N-terminal regions in the two nsp8 subunits (**Figs. 2, 3**). These prominent nsp8 extensions reach up to 26 base pairs away from the active site and use positively charged residues to form multiple RNA backbone interactions (**Fig. 3**). The two nsp8 extensions form different RNA interactions, arguing for sequence-independent binding. The nsp8 extensions get ordered along RNA, as they are flexible in nsp8-nsp7 complexes^18,19^.

The interactions of the nsp8 extensions with exiting RNA may explain the processivity of the RdRp, which is required for the replication of the very long RNA genome of coronaviruses and other viruses of the nidovirus family^5^. It is known that nsp8 and nsp7 confer processivity to nsp12, and that mutation of residue K58 is lethal for the virus^15^. K58 is located in the nsp8 extension, and interacts with exiting RNA (**Fig. 3c**). Thus the nsp8 extensions may be regarded as ‘sliding poles’ that slide along exiting RNA at the rear of the polymerase to prevent premature dissociation of the RdRp during replication. The sliding poles could serve a function similar to the ‘sliding clamps’ of DNA replication machineries^23^.

To investigate how the RdRp binds the incoming nucleoside triphosphate (NTP), we superimposed our structure onto the related structure of the norovirus RdRp-nucleic acid complex^22^. This suggested that contacts between nsp12 and the NTP are conserved (**Extended Data Fig. 3c**). The nsp12 residue N691 may recognize the 2’-OH group of the NTP, thereby rendering the RdRp specific for the synthesis of RNA, rather than DNA. Our modeling is also consistent with binding of the triphosphorylated form of remdesivir to the NTP site, because there is space in the NTP site to accommodate the additional nitrile group that is present at the 1’ position of the ribose ring in this nucleoside analogue.

When our study was about to be completed, a manuscript became available that also describes a structure of a SARS-CoV-2 RdRp-RNA complex^24^. Whereas the core structures appear to be very similar, we additionally observe exiting RNA and novel nsp8 extensions that are implicated in enzyme processivity. The other study suggested that remdesivir functions as an immediate RNA chain terminator^24^. However, this contradicts previous biochemistry^10,25^ that showed that remdesivir causes delayed chain termination after the addition of several more nucleotides. To resolve this, we will study the mechanism of RdRp inhibition by remdesivir with a combination of biochemistry and structural biology in the future.

## EXPERIMENTAL PROCEDURES

No statistical methods were used to predetermine sample size. The experiments were not randomized, and the investigators were not blinded to allocation during experiments and outcome assessment.

### Cloning and protein expression

The SARS-CoV-2 nsp12 gene was codon optimized for expression in insect cells. The SARS-CoV-2 nsp8 and nsp7 genes were codon optimized for expression in Escherichia coli. Synthesis of genes was performed by GeneArt (ThermoFischer Scientific GENEART GmbH, Regensburg, Deutschland). The gene synthesis products of the respective genes were PCR amplified with ligation-independent cloning (LIC) compatible primer pairs (nsp12: Forward primer: 5’-TACTTC CAA TCC AAT GCA TCT GCT GAC GCT CAG TCC TTC CTG-3’, reverse primer: 5’-TTA TCC ACT TCC AAT GTT ATT ATT GCA GCA CGG TGT GAG GGG-3’; nsp8: Forward primer: 5’-TAC TTC CAA TCC AAT GCA GCA ATT GCA AGC GAA TTT AGC AGC CTG-3’, reverse primer: 5’-TTA TCC ACT TCC AAT GTT ATT ACT GCA GTT TAA CTG CGC TAT TTG CAC G-3’; nsp7: Forward primer: 5’-TAC TTC CAA TCC AAT GCA AGC AAA ATG TCC GAT GTT AAA TGC ACC AGC-3, reverse primer: 5’-TTA TCC ACT TCC AAT GTT ATT ACT GCA GGG TTG CAC GAT TAT CCA GC-3’). The PCR products for nsp8 and nsp7 were individually cloned into the pET derived vector 14-B (a gift from S. Gradia, UC Berkeley, Ad-dgene: 48308). The two constructs for nsp8 and nsp7 contain an N-terminal 6xHis tag and a tobacco etch virus protease cleavage site. The PCR product containing codon optimized nsp12 was cloned into the modified pFastBac vector 438-C (a gift from S. Gradia, UC Berkeley, Addgene: 55220) via LIC. The nsp12 construct contained an N-terminal 6xHis tag, followed by an MBP tag, a 10xAsp sequence, and a tobacco etch virus protease cleavage site. All constructs were verified by sequencing.

The SARS-CoV-2 nsp12 plasmid (500 ng) was transformed into DH10EMBacY cells using electroporation to generate a bacmid encoding full-length nsp12. Virus production and expression in insect cells was then performed as described^26^. After 60 hours of expression in Hi5 cells, cells were collected by centrifugation and resuspended in lysis buffer (300 mM NaCl, 50 mM Na-HEPES pH 7.4, 10 % (v/v) glycerol, 30 mM imidazole pH 8.0, 3 mM MgCl2, 5 mM β-mercaptoethanol, 0.284 μg ml-1 leupeptin, 1.37 μg ml-1 pepstatin, 0.17 mg ml-1 PMSF, and 0.33 mg ml-1 benzamidine). The SARS-CoV-2 nsp8 and nsp7 plasmids were overexpressed in E. coli BL21 (DE3) RIL cells grown in LB medium. Cells were grown to an OD of 600 at 37 °C and protein expression was subsequently induced with 0. 5 mM isopropyl β-D-1-thiogalactopyranoside at 18 °C for 16 hours. Cells were collected by centrifugation and resuspended in lysis buffer (300 mM NaCl, 50 mM Na-HEPES pH 7.4, 10 % (v/v) glycerol, 30 mM imidazole pH 8.0, 5 mM β-mercaptoeth-anol, 0.284 μg ml-1 leupeptin, 1.37 μg ml-1 pepstatin, 0.17 mg ml-1 PMSF, and 0.33 mg ml-1 benzamidine).

### Protein purification

Protein purifications were performed at 4 °C. After harvest and resuspension, cells of the SARS-CoV-2 nsp12 expression were immediately sonicated for cell lysis. Lysates were subsequently cleared by centrifugation (87,207g, 4 °C, 30 min) and ultracentrifugation (235,000g, 4 °C, 60 min). The supernatant containing nsp12 was filtered using a 5-μm syringe filter, followed by filtration with a 0.8-μm syringe filter (Millipore) and applied onto a HisTrap HP 5 mL (GE Healthcare), preequilibrated in lysis buffer (300 mM NaCl, 50 mM Na-HEPES pH 7.4, 10 % (v/v) glycerol, 30 mM imidazole pH 8.0, 3 mM MgCl2, 5 mM β-mercaptoethanol, 0.284 μg ml-1 leupeptin, 1.37 μg ml-1 pepstatin, 0.17 mg ml-1 PMSF, and 0.33 mg ml-1 benzamidine). After application of the sample, the column was washed with 6 CV high salt buffer (1000 mM NaCl, 50 mM Na-HEPES pH 7.4, 10 % (v/v) glycerol, 30 mM imidazole pH 8.0, 3 mM MgCl2, 5 mM β-mercaptoethanol, 0.284 μg ml-1 leupeptin, 1.37 μg ml-1 pepstatin, 0.17 mg ml-1 PMSF, and 0.33 mg ml-1 benzamidine), and 6 CV lysis buffer. The HisTrap was then attached to an XK column 16/20 (GE Healthcare), prepacked with amylose resin (New England Biolabs), which was pre-equilibrated in lysis buffer. The protein was eluted from the HisTrap column directly onto the amylose column using nickel elution buffer (300 mM NaCl, 50 mM Na-HEPES pH 7.4, 10 % (v/v) glycerol, 500 mM imidazole pH 8.0, 3 mM MgCl2, 5 mM β-mercaptoethanol). The HisTrap column was then removed and the amylose column was washed with 10 CV of lysis buffer. Protein was eluted from the amylose column using amylose elution buffer (300 mM NaCl, 50 mM Na-HEPES pH 7.4, 10 % (v/v) glycerol, 116.9 mM maltose, 30 mM imidazole pH 8.0, 5 mM β-mercaptoethanol). Peak fractions were assessed via SDS-PAGE and staining with Coomassie. Peak fractions containing nsp12 were pooled and mixed with 8 mg of His-tagged TEV protease. After 12 hours of protease digestion, protein was applied to a HisTrap column equilibrated in lysis buffer to remove uncleaved nsp12, His6-MBP, and TEV. Subsequently, the flow-through containing nsp12 was applied to a HiTrap Heparin 5 column mL (GE Healthcare). The flow-through containing nsp12 was collected and concentrated in a MWCO 50,000 Amicon Ultra Centrifugal Filter unit (Merck). The concentrated sample was applied to a HiLoad S200 16/600 pg equilibrated in size exclusion buffer (300 mM NaCl, 20 mM Na-HEPES pH 7.4, 10 % (v/v) glycerol, 1 mM MgCl2, 1 mM TCEP). Peak fractions were assessed by SDS-PAGE and Coomassie staining. Peak fractions were pooled and concentrated in a MWCO 50,000 Amicon Ultra Centrifugal Filter (Merck). The concentrated protein with a final concentration of 102 μM was aliquoted, flash-frozen, and stored at −80 °C until use.

SARS-CoV-nsp8 and nsp7 were purified separately using the same purification procedure, as follows. After cell harvest and resuspension in lysis buffer, the protein of interest was immediately sonicated. Lysates were subsequently cleared by centrifugation (87.200g, 4 °C, 30 min). The supernatant was applied to a HisTrap HP 5 mL column (GE Healthcare), preequilibrated in lysis buffer. The column was washed with 9.5 CV high salt buffer (1000 mM NaCl, 50 mM Na-HEPES pH 7.4, 10 % (v/v) glycerol, 30 mM imidazole pH 8.0, 5 mM β-mercaptoethanol, 0.284 μg ml-1 leupeptin, 1.37 μg ml-1 pepstatin, 0.17 mg ml-1 PMSF, and 0.33 mg ml-1 benzamidine), and 9.5 CV low salt buffer (150 mM NaCl, 50 mM Na-HEPES pH 7.4, 10 % (v/v) glycerol, 30 mM imidazole pH 8.0, 5 mM β-mercaptoethanol). The sample was then eluted using nickel elution buffer (150 mM NaCl, 50 mM Na-HEPES pH 7.4, 10 % (v/v) glycerol, 500 mM imidazole pH 8.0, 5 mM β-mercaptoethanol). The eluted protein was dialyzed in dialysis buffer (150 mM NaCl, 50 mM Na-HEPES pH 7.4, 10 % (v/v) glycerol, 5 mM β-mercaptoethanol) in the presence of 2 mg His-tagged TEV protease. After 12 hours, imidazole pH 8.0 was added to a final concentration of 30 mM. The dialyzed sample was subsequently applied to a HisTrap HP 5 mL column (GE Healthcare), preequilibrated in dialysis buffer. The flow-through that contained the protein of interest was then applied to a HiTrap Q 5 mL column (GE Healthcare). The Q column flow-through containing nsp8 or nsp7 was concentrated using a MWCO 3 kDa Amicon Ultra Centrifugal Filter (Merck) and applied to a HiLoad S200 16/600 pg equilibrated in size exclusion buffer (150 mM NaCl, 20 mM Na-HEPES pH 7.4, 5 % (v/v) glycerol, 1 mM TCEP). Peak fractions were assessed by SDS-PAGE and Coomassie staining. Peak fractions were pooled. Nsp7 with a final concentration of 418 μM was aliquoted, flash-frozen, and stored at −80 °C until use. Nsp8 with a final concentration of 250 μM was aliquoted, flash-frozen, and stored at −80 °C until use. All protein identities were confirmed by mass spectrometry.

### RNA extension assays

All RNA oligos were purchased from Integrated DNA Technologies. The RNA sequence used for the transcription assay is /56-FAM/rUrUrU rUrCrA rUrGrC rUrArC rGrCrG rUrArG rUrUr UrUrC rUrArC rGrCrG. We designed a minimal substrate by connecting the template RNA to the RNA primer by a tetraloop, to protect the blunt ends of the RNA duplex and to ensure efficient annealing. RNA was annealed in 50 mM NaCl and 10 mM Na-HEPES pH 7.5 by heating the solution to 75 °C and gradually cooling to 4 °C. RNA extension reactions contained RNA (5 μM), nsp12 (5 μM), nsp8 (15 μM) and nsp7 (15 μM) in 100 mM NaCl, 20 mM Na-HEPES pH 7.5, 5 % (v/v) glycerol, 10 mM MgCl2 and 5 mM β-mercaptoethanol. Reactions were incubated at 37 °C for 5 min and the RNA extension was initiated by addition of NTPs (150 μM UTP, GTP and CTP, and 300 μM ATP). Reactions were stopped by the addition of 2X stop buffer (7M urea, 50 mM EDTA pH 8.0, 1x TBE buffer). Samples were digested with proteinase K (New England Biolabs) and RNA products were separated on 20% acrylamide gels in 1X TBE buffer supplemented with 8M urea. 6-FAM labeled RNA products were visualized by Typhoon 95000 FLA Imager (GE Healthcare Life Sciences).

### Cryo-EM sample preparation and data collection

An RNA scaffold for RdRP-RNA complex formation was annealed by mixing equimolar amounts of two RNA strands (5’-rUrUrU rUrCrA rUrGrC rUrArC rGrCrG rUrArG-3’; 56-FAM/rCrUrA rCrGrC rG-3’) (IDT Technologies) in annealing buffer (10 mM Na-HEPES pH 7.4, 50 mM NaCl) and heating to 75 °C, followed by step-wise cooling to 4 °C. For complex formation, 1.2 nmol of purified nsp12 was mixed with a 1.2-fold molar excess of RNA scaffold and 6-fold molar excess of each nsp8 and nsp7. After incubation at room temperature for 10 min, the EC was subjected to size exclusion chromatography on a Superdex 200 Increase 3.2/300 equilibrated with complex buffer (20 mM Na-HEPES pH 7.4, 100 mM NaCl, 1 mM MgCl2, 1 mM TCEP). Peak fractions corresponding to a nucleic-acid rich high-molecular weight population (as judged by absorbance at 260 nm) were pooled and concentrated in a MWCO 30,000 Vivaspin 500 concentrator (Sartorius) to approx. 20 μl. 3 μL of the concentrated RdRp-RNA complex were mixed with 0.5 μl of octyl ß-D-glucopyranoside (0.003% final concentration) and applied to freshly glow discharged R 2/1 holey carbon grids (Quantifoil). Prior to flash freezing in liquid ethane, the grid was blotted for 6 seconds with a blot force of 5 using a Vitrobot MarkIV (Thermo Fischer Scientific) at 4°C and 100% humidity.

Cryo-EM data collection was performed with SerialEM^27^ using a Titan Krios transmission electron microscope (Thermo Fischer Scientific) operated at 300 keV. Images were acquired in EFTEM mode with a slit with of 20 eV using a GIF quantum energy filter and a K3 direct electron detector (Gatan) at a nominal magnification of 105,000x corresponding to a calibrated pixel size of 0.834 Å/pixel. Exposures were recorded in counting mode for 2.2 seconds with a dose rate of 19 e-/px/s resulting in a total dose of 60 e-/Å2 that was fractionated into 80 movie frames. Because initial processing showed that the particles adopted only a limited number of orientations in the vitreous ice layer, a total of 8168 movies were collected at 30° stage tilt to yield a broader distribution of orientations. Untilted data was discarded. Motion correction, CTF-estimation, and particle picking and extraction were performed on the fly using Warp^28^.

### Cryo-EM data processing and analysis

1.3 million particles were exported from Warp^28^ to cryoSPARC^29^, and the particles were subjected to 2D classification. 25% of the particles were selected from classes deemed to represent the polymerase, and refined against a synthetic reference prepared from PDB-6M71. Ab initio refinement was performed using particles from bad 2D classes to obtain five 3D classes of ‘junk’. These five classes and the first polymerase reconstruction were used as starting references to sort the initial 1.3M particles in supervised 3D classification rather than 2D, as the latter tended to exclude less abundant projection directions. 514k particles (39%) from the resulting polymerase class were subjected to another ab initio refinement to obtain five starting references containing four ‘junk’ classes and the complex of interest. These classes were used as starting references in another supervised 3D classification. 418k particles (82%) from the class representing the complex were exported from cryoSPARC to RELION 3.0^30^. There, all particles were refined in 3D against the reconstruction previously obtained in cryoSPARC, using a mask including only the core part of the polymerase and a short segment of upstream RNA to obtain a 3.1 Å reconstruction. CTF refinement and another round of 3D refinement improved the resolution further to 2.9 Å. Particles were re-extracted at 1.3 Å/ px in a bigger box in Warp to accommodate distant parts of the RNA. Unsupervised 3D classification with local alignment was performed to obtain 2 classes: with nsp8b present, and without. 172k particles with nsp8b present were finally subjected to focused 3D refinement using a mask including the RNA, nsp8a and nsp8b.

### Model building and refinement

To build the atomic model of the RdRp-RNA complex, we started from the structure of the free SARS-CoV-2 RdRp (PDB: 6M71) that was recently slightly adjusted by Tristan Croll (Cambridge University, UK; https://twitter.com/CrollTristan/status/1247846163061133312). The structure was rigid-body fit into the cryo-EM reconstruction and adjusted manually in Coot^31^. After adjustment of the protein subunits, unmodeled density remained for helical segments in the N-terminal regions of both copies of nsp8. These nsp8 extensions were modeled by superimposing the nsp8 model (PDB: 2AHM; chain H) from the crystal structure of the nsp7-nsp8 hexadecamer^18^, in which the far N-terminal region of nsp8 adopts a similar fold. Careful inspection of the remaining A-form RNA density revealed that in our complex, instead of the originally designed short template-primer duplex (see above), four copies of one of the RNA strands were annealed to form a pseudo-continuous, longer RNA duplex. Annealing was mediated by a 10 bp self-complementary region within this RNA strand (Extended Date Figure 1c). Nucleotides 5-18 of four RNA strands were modeled, whereas the flapped-out nucleotides 1-4 were invisible due to mobility and excluded from the model. The model was real-space refined using phenix.refine^32^ against a composite map of the focused refinement and global reconstructions generated in phenix.combine_focused_maps and shows excellent stereochemistry (Extended Data Table 1). Figures were prepared with PyMol and Chimera^33^.

## ACKNOWLEDGEMENTS

We thank all members of the Department of Molecular Biology at the Max-Planck-Institute for Biophysical Chemistry for support. We thank Jana Schmitzova for helpful discussions and advice. H.S.H. was supported by the Deutsche Forschungs-gemeinschaft (FOR2848). P.C. was supported by the Deutsche Forschungsgemeinschaft (EXC 2067/1 39072994, SFB860, SPP2191), the ERC Advanced Investigator Grant TRANSREG-ULON (grant agreement No 693023) and the Volkswagen Foundation. We thank Henning Urlaub (Max Planck Institute for Biophysical Chemistry) for protein identification by mass spectrometry.

## AUTHOR CONTRIBUTIONS

H.S.H., G.K., L.F., C.D. and D.T. designed and carried out experiments and data analysis. P.C. designed and supervised research. All authors interpreted data and wrote the manuscript.

## DATA AVAILABILITY

Reconstructions and structure coordinate files will be made available in the Electron Microscopy Database and the Protein Data Bank.

---

**Extended Data Figure 1.**
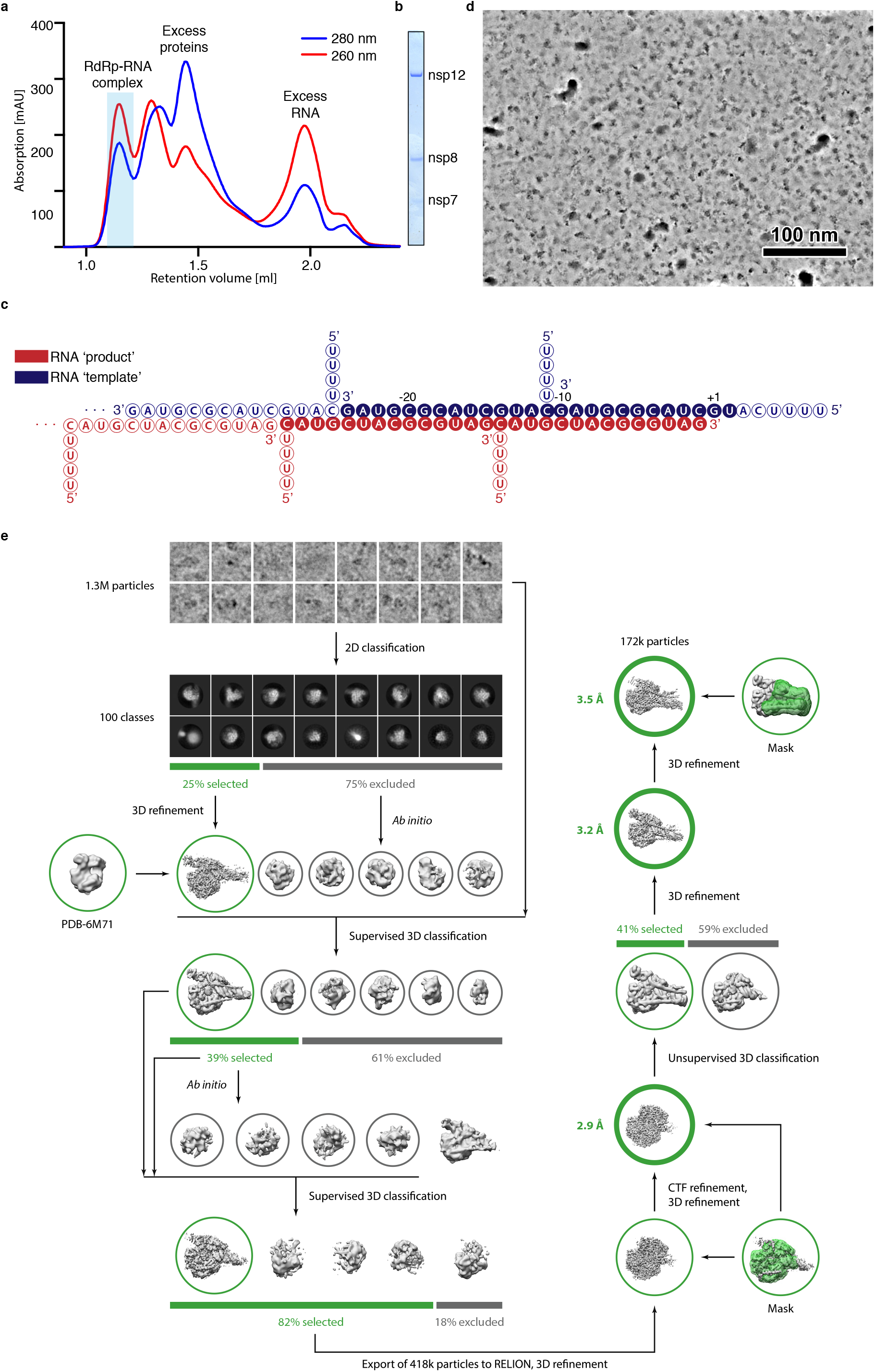
Cryo-EM analysis. Related to Figures 1, 2. **a** Purification of RdRp-RNA complex by size exclusion chromatography. The peak used for structural analysis is highlighted in blue. **b** Exemplary SDS-PAGE analysis of purified complex with RdRp subunits labeled. **c** RNA duplex scaffold formed by oligomerization of a short pseudo-palindromic RNA. The depicted base pairing gave rise to a pseudo-continuous A-form duplex. Solid and hollow circles show RNA nucleotides that were included in the structure or not visible, respectively. **d** Example denoised micrograph. Scale bar, 100 nm. **e** Cryo-EM processing tree.

**Extended Data Figure 2.**
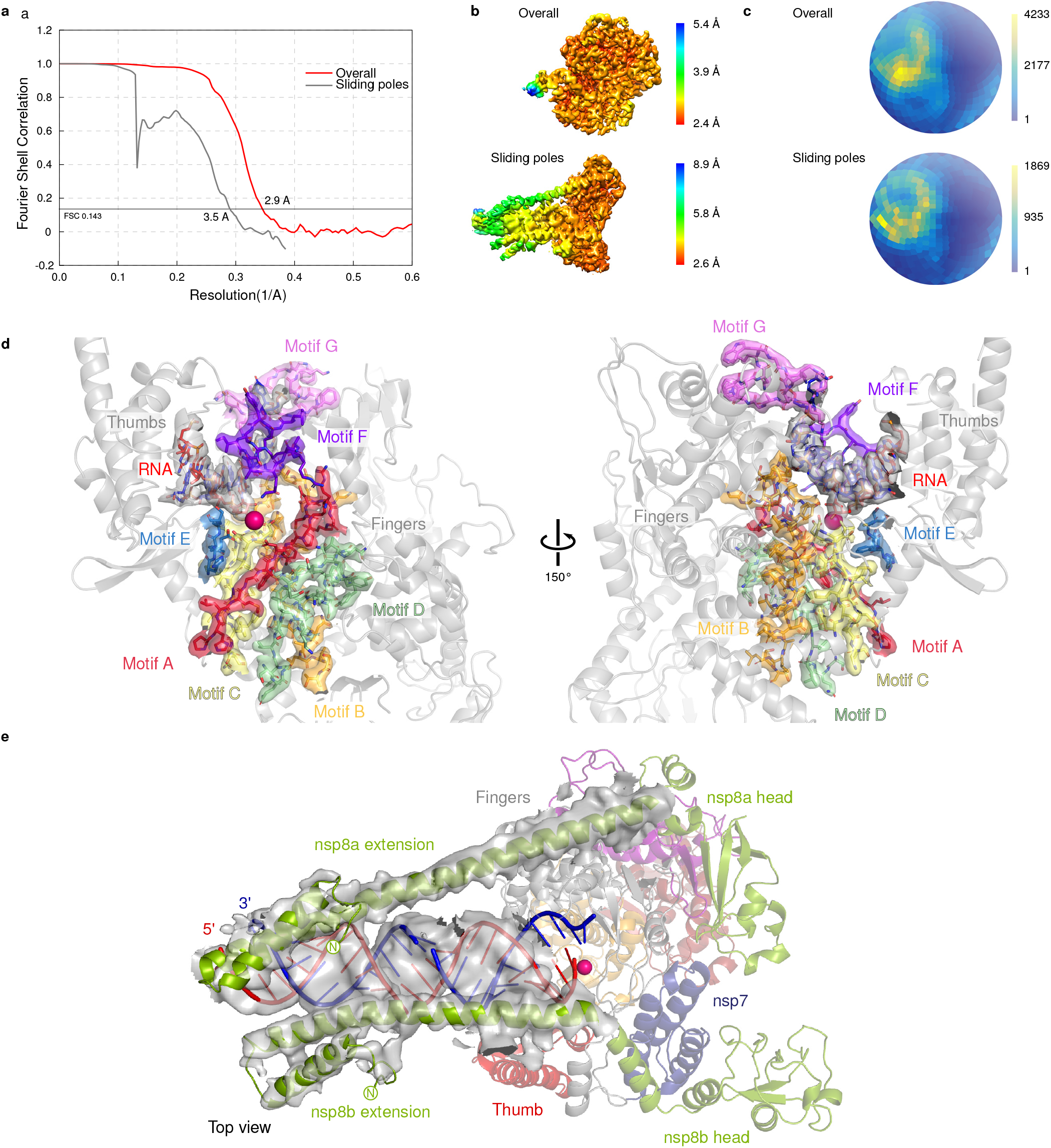
Cryo-EM reconstructions. Related to Figure 2. **a** Fourier shell correlation (FSC) plots for reported reconstructions and resolution estimation. **b** Local resolution distribution. **c** Angular distribution plots. Scale shows the number of particles assigned to a particular angular bin. Blue, a low number of particles; yellow, a high number of particles. **d** Cryo-EM map for the RdRp active center region including elements with sequence motifs A-G. The active site is indicated by a magenta sphere. **e** Cryo-EM map for the RNA duplex and the nsp8 extensions. The active site is indicated by a magenta sphere.

**Extended Data Figure 3.**
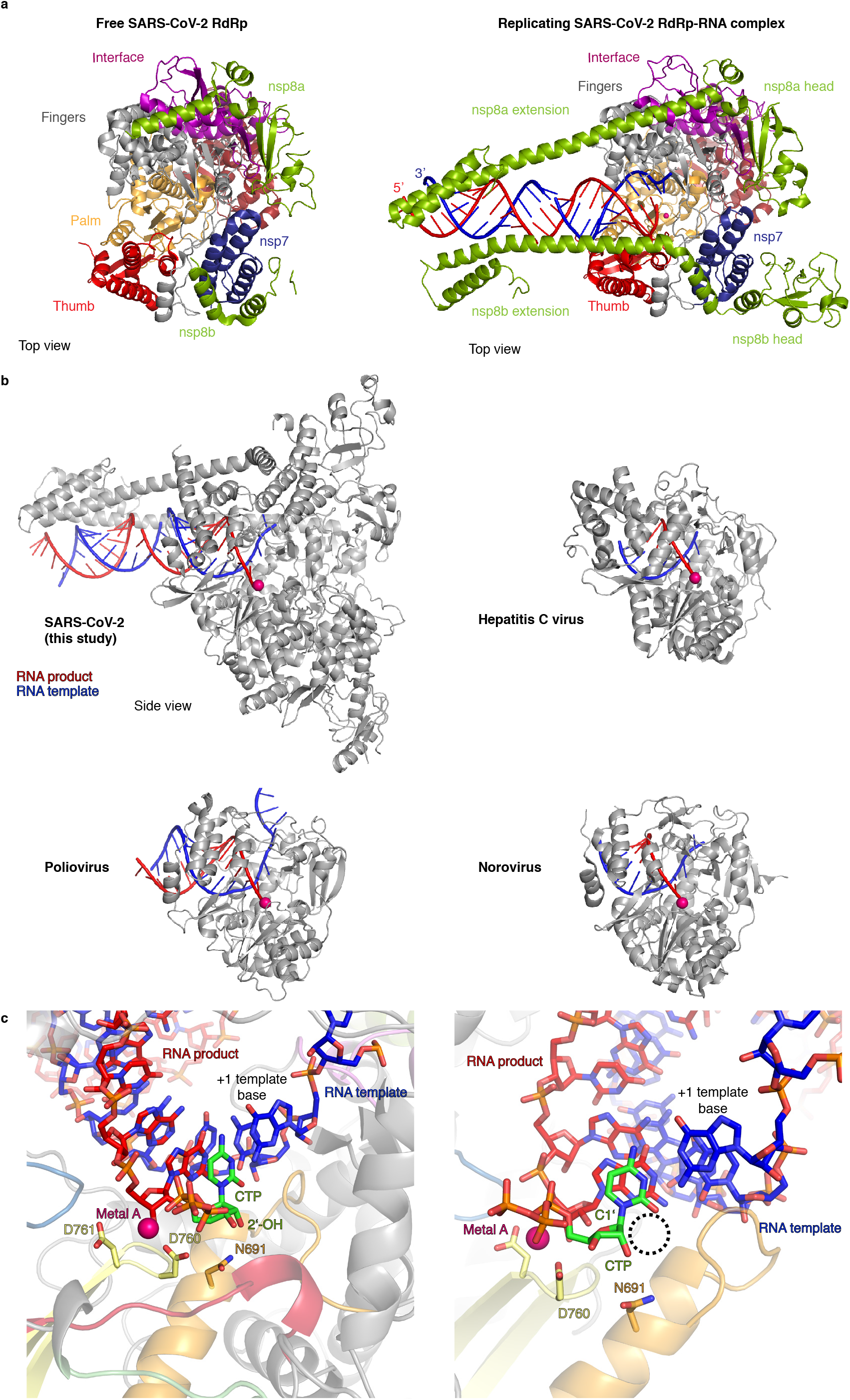
Structural comparisons. Related to Figures 2, 3. **a** Comparison of the free SARS-CoV-2 RdRp structure16 (left) and the replicating RdRp-RNA complex (right, this study). Color code as in Fig. 2. **b** Similar location and orientation of the RNA template-product duplex in RdRp complexes of SARS-CoV-2 virus (top left, this study), hepatitis C virus20 (top right), poliovirus21 (bottom left), and norovirus22 (bottom right). Structures are shown as ribbon models with RNA template and product strands in blue and red, respectively. An active site metal ion is shown as a magenta sphere. Side view as defined in Fig. 2. **c** Model of substrate nucleoside triphosphate (NTP) in the RdRp active site. A CTP substrate was placed after superposition of the norovirus RdRp-nucleic acid complex structure22. Coloring as in Fig. 3b. Active site residues D760, D761 and N691 are shown as sticks, and the modeled active site metal ion is shown as a magenta sphere. When the nucleoside triphosphate form of remdesivir would bind in the NTP site, the nitrile group connected to the ribose C1’ position would be accommodated in the space indicated by the dashed circle.

## EXTENDED DATA TABLE

**ExtendedData Table 1.**
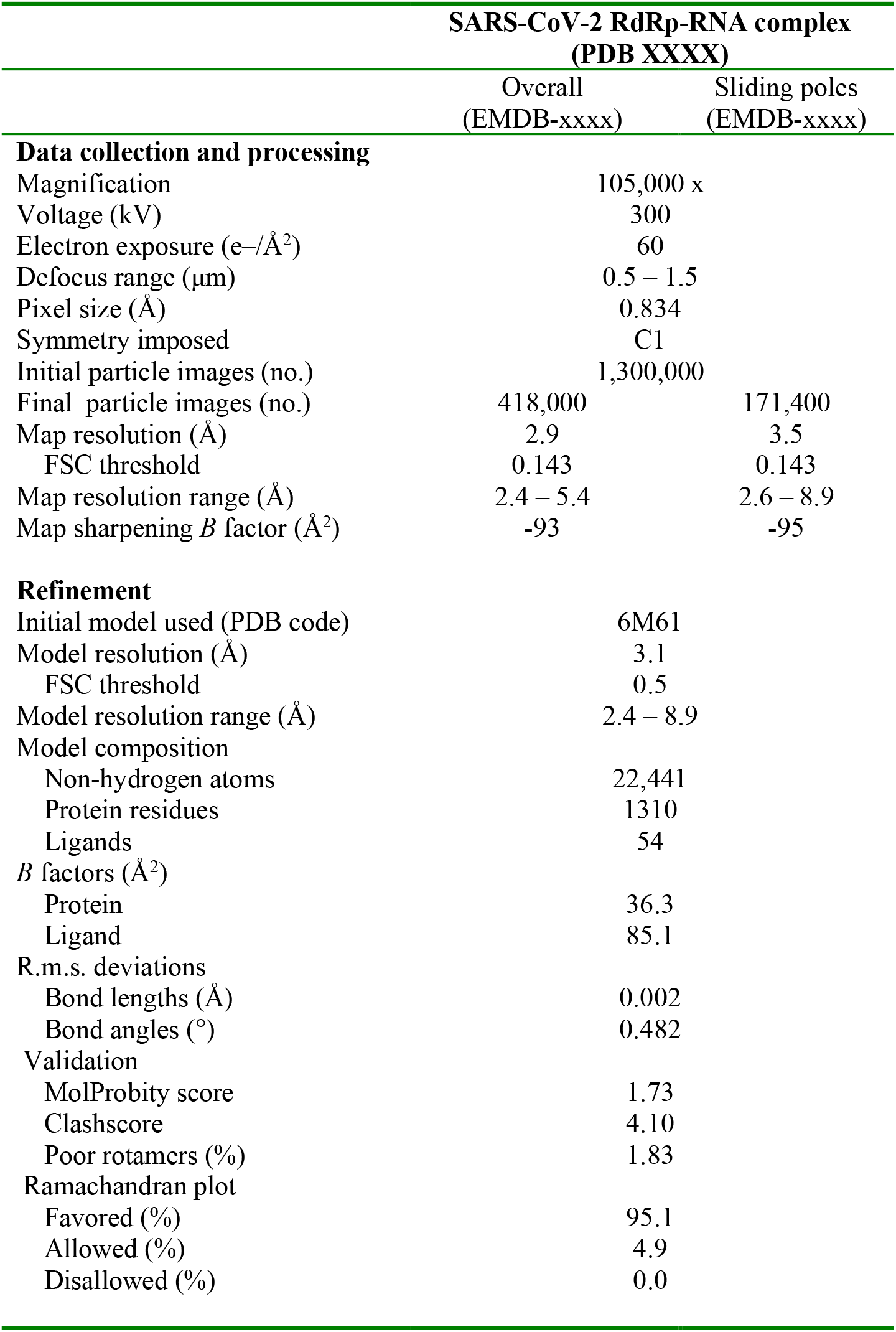
Cryo-EMdatacollection andprocessinginformation.

## REFERENCES

1. Hilgenfeld, R. & Peiris, M. From SARS to MERS: 10 years of research on highly pathogenic human coronaviruses. Antiviral Res 100, 286–295, doi:10.1016/j.antiviral.2013.08.015 (2013).

2. Wu, F. et al. A new coronavirus associated with human respiratory disease in China. Nature 579, 265–269, doi:10.1038/s41586-020-2008-3 (2020).

3. Zhou, P. et al. A pneumonia outbreak associated with a new coronavirus of probable bat origin. Nature 579, 270–273, doi:10.1038/s41586-020-2012-7 (2020).

4. Snijder, E. J., Decroly, E. & Ziebuhr, J. The Nonstructural Proteins Directing Coronavirus RNA Synthesis and Processing. Adv Virus Res 96, 59–126, doi:10.1016/bs.aivir.2016.08.008 (2016).

5. Posthuma, C. C., Te Velthuis, A. J. W & Snijder, E. J. Nidovirus RNA polymerases: Complex enzymes handling exceptional RNA genomes. Virus Res 234, 58–73, doi:10.1016/j.virusres.2017.01.023 (2017).

6. Subissi, L. et al. SARS-CoV ORF1b-encoded nonstructural proteins 12-16: replicative enzymes as antiviral targets. Antiviral Res 101, 122–130, doi:10.1016/j.antiviral.2013.11.006 (2014).

7. Jordan, P. C., Stevens, S. K. & Deval, J. Nucleosides for the treatment of respiratory RNA virus infections. Antivir Chem Chemother 26, 2040206618764483, doi:10.1177/2040206618764483 (2018).

8. Sheahan, T. P. et al. Broad-spectrum antiviral GS-5734 inhibits both epidemic and zoonotic coronaviruses. Sci Transl Med 9, doi:10.1126/scitranslmed.aal3653 (2017).

9. Agostini, M. L. et al. Coronavirus Susceptibility to the Antiviral Remdesivir (GS-5734) Is Mediated by the Viral Polymerase and the Proofreading Exoribonuclease. mBio 9, doi:10.1128/mBio.00221-18 (2018).

10. Gordon, C. J., Tchesnokov, E. P., Feng, J. Y., Porter, D. P & Gotte, M. The antiviral compound remdesivir potently inhibits RNA-dependent RNA polymerase from Middle East respiratory syndrome coronavirus. J Biol Chem, doi:10.1074/jbc.AC120.013056 (2020).

11. Kupferschmidt, K. & Cohen, J. Race to find COVID-19 treatments accelerates. Science 367, 1412–1413, doi:10.1126/science.367.6485.1412 (2020).

12. Ko, W. C. et al. Arguments in favour of remdesivir for treating SARS-CoV-2 infections. Int J Antimicrob Agents, 105933, doi:10.1016/j.ijantimicag.2020.105933 (2020).

13. Cao, Y. C., Deng, Q. X. & Dai, S. X. Remdesivir for severe acute respiratory syndrome coronavirus 2 causing COVID-19: An evaluation of the evidence. Travel Med Infect Dis, 101647, doi:10.1016/j.tmaid.2020.101647 (2020).

14. Ahn, D. G., Choi, J. K., Taylor, D. R. & Oh, J. W. Biochemical characterization of a recombinant SARS coronavirus nsp12 RNA-dependent RNA polymerase capable of copying viral RNA templates. Arch Virol 157, 2095–2104, doi:10.1007/s00705-012-1404-x (2012).

15. Subissi, L. et al. One severe acute respiratory syndrome coronavirus protein complex integrates processive RNA polymerase and exonuclease activities. Proc Natl Acad Sci U S A 111, E3900–3909, doi:10.1073/pnas.1323705111 (2014).

16. Gao, Y. et al. Structure of the RNA-dependent RNA polymerase from COVID-19 virus. Science, eabb7498, doi:10.1126/science.abb7498 (2020).

17. Kirchdoerfer, R. N. & Ward, A. B. Structure of the SARS-CoV nsp12 polymerase bound to nsp7 and nsp8 co-factors. Nat Commun 10, 2342, doi:10.1038/s41467-019-10280-3 (2019).

18. Zhai, Y. et al. Insights into SARS-CoV transcription and replication from the structure of the nsp7-nsp8 hexadecamer. Nat Struct Mol Biol 12, 980–986, doi:10.1038/nsmb999 (2005).

19. Xiao, Y. et al. Nonstructural proteins 7 and 8 of feline coronavirus form a 2:1 heterotrimer that exhibits primer-independent RNA polymerase activity. J Virol 86, 4444–4454, doi:10.1128/JVI.06635-11 (2012).

20. Appleby, T. C. et al. Viral replication. Structural basis for RNA replication by the hepatitis C virus polymerase. Science 347, 771–775, doi:10.1126/science.1259210 (2015).

21. Gong, P. & Peersen, O. B. Structural basis for active site closure by the poliovirus RNA-dependent RNA polymerase. Proc Natl Acad Sci U S A 107, 22505–22510, doi:10.1073/pnas.1007626107 (2010).

22. Zamyatkin, D. F., Parra, F., Machin, A., Grochulski, P. & Ng, K. K. Binding of 2’-amino-2’-deoxycytidine-5’-triphosphate to norovirus polymerase induces rearrangement of the active site. J Mol Biol 390, 10–16, doi:10.1016/j.jmb.2009.04.069 (2009).

23. Moldovan, G. L., Pfander, B. & Jentsch, S. PCNA, the maestro of the replication fork. Cell 129, 665–679, doi:10.1016/j.cell.2007.05.003 (2007).

24. Yin, W. et al. Structural Basis for the Inhibition of the RNA-Dependent RNA Polymerase from SARS-CoV-2 by Remdesivir. bioRxiv, 2020.2004.2008.032763, doi:10.1101/2020.04.08.032763 (2020).

25. Tchesnokov, E. P., Feng, J. Y., Porter, D. P. & Gotte, M. Mechanism of Inhibition of Ebola Virus RNA-Dependent RNA Polymerase by Remdesivir. Viruses 11, doi:10.3390/v11040326 (2019).

26. Vos, S. M. et al. Architecture and RNA binding of the human negative elongation factor. Elife 5, doi:10.7554/eLife.14981 (2016).

27. Mastronarde, D. N. Automated electron microscope tomography using robust prediction of specimen movements. J Struct Biol 152, 36–51, doi:10.1016/j.jsb.2005.07.007 (2005).

28. Tegunov, D. & Cramer, P. Real-time cryo-electron microscopy data preprocessing with Warp. Nat Methods 16, 1146–1152, doi:10.1038/s41592-019-0580-y (2019).

29. Punjani, A., Rubinstein, J. L., Fleet, D. J. & Brubaker, M. A. cryo-SPARC: algorithms for rapid unsupervised cryo-EM structure determination. Nat Methods 14, 290–296, doi:10.1038/nmeth.4169 (2017).

30. Zivanov, J., Nakane, T. & Scheres, S. H. W. A Bayesian approach to beam-induced motion correction in cryo-EM single-particle analysis. IUCrJ 6, 5–17, doi:10.1107/S205225251801463X (2019).

31. Emsley, P., Lohkamp, B., Scott, W. G. & Cowtan, K. Features and development of Coot. Acta Crystallogr D Biol Crystallogr 66, 486–501, doi:10.1107/S0907444910007493 (2010).

32. Afonine, P. V. et al. Real-space refinement in PHENIX for cryo-EM and crystallography. Acta Crystallogr D Struct Biol 74, 531–544, doi:10.1107/S2059798318006551 (2018).

33. Huang, C. C., Meng, E. C., Morris, J. H., Pettersen, E. F. & Ferrin, T. E. Enhancing UCSF Chimera through web services. Nucleic Acids Res 42, W478–484, doi:10.1093/nar/gku377 (2014).

